# Unraveling membrane protein localization and stabilization in nanodiscs

**DOI:** 10.1101/2023.07.20.549795

**Authors:** So-Jung Kim, Young Hoon Koh, Soung-Hun Roh

## Abstract

Nanodiscs are nanoscale structures consisting of a lipid bilayer surrounded by membrane scaffold proteins (MSPs). They are widely used in the study of membrane proteins (MPs) because they provide a stable lipid environment. However, the precise mechanism governing MP behavior within the nanodisc remains elusive. Here, we examined the cryo-EM structures of various MPs reconstituted in nanodiscs from an electron microscopy database (EMPIAR). By analyzing the heterogeneity and interactions in the nanodiscs, we found that MPs within nanodiscs display a distinct spatial preference toward the edges of the nanodisc shells. Furthermore, we observed that MPs can establish direct, amphipathic interactions with the MSPs, promoting protein stability. These interactions may induce a rearrangement of the MSP-MSP interactions, leading to the formation of MP-MSP interactions Collectively, our study provides structural and biophysical insights into how nanodiscs contribute to MP structural behavior and stability.

**SIGNIFICANCE:** By thoroughly examining multiple deposited datasets of membrane proteins (MPs) reconstituted in nanodiscs, we have gathered compelling evidence that MPs exhibit a clear spatial inclination toward the periphery of the nanodisc shells. Moreover, we have observed that MPs establish direct and amphipathic interactions with membrane scaffold proteins (MSPs). These interactions have the potential to induce a rearrangement of the MSP-MSP interactions, consequently forming MP-MSP interactions. Through quantitative analysis, we have successfully characterized the significant role played by these interactions in ensuring the overall stability of the proteins.

## INTRODUCTION

Membrane proteins (MPs) are essential for various biological processes, such as transport, enzyme activity, and signal transduction <u>(Sligar and Denisov 2021)</u>. However, the preparation of MPs poses significant challenges, including aggregation, loss of function, and difficulty in crystallization. Recent advancements in cryo-EM and various MP reconstitution methods have overcome these limitations <u>(Januliene and Moeller 2021)</u>. Two prominent approaches for solubilizing MPs use detergents and nanodiscs <u>(Zampieri et al. 2021)</u>. While the structural determination of MPs is commonly performed using samples in detergent, concerns exist for potential differences between the detergent and native environments. Nanodiscs are architecturally based on high-density lipoproteins surrounded by apolipoprotein A1, an amphipathic helical protein <u>(Patel et al. 2019)</u>. The engineered apolipoprotein A1 functions as a membrane scaffold protein (MSP) and envelops the hydrophobic transmembrane (TM) regions of the MPs embedded within the lipid bilayer. The nanodisc environment provides a lipidic bilayer environment closely mimicking the native state of the MP and maintaining its stable conformation during the purification process. Moreover, several reports analyzing the structures of TRPV1 <u>(Gao et al. 2016)</u> and NompC <u>(Gao et al. 2016; Autzen et al. 2019)</u> suggest that reconstitution in nanodiscs enables the determination of higher resolution structures through cryo-EM.

Despite the increasing recognition of the benefits of nanodiscs in diverse biophysical approaches, there is a lack of comprehensive experimental data on the properties of nanodiscs when used with embedded MPs. This study examined MP structures reconstituted in nanodiscs in the Electron Microscopy Public Image Archive (EMPIAR) <u>(Iudin</u> <u>et al. 2016)</u>. Our analysis focused on the structural heterogeneity in nanodiscs, which revealed that nanodiscs possess structural fluidity and that MPs move laterally within the nanodiscs. This behavior has been observed previously *in silico* <u>(López et al. 2019</u>; <u>Orioli et al. 2021</u>). Kern et al. also experimentally observed interactions between SARS-CoV-2 ORF3a and MSP in the nanodisc environment <u>(Kern et al. 2021)</u>. The study provides further evidence of such MP-MSP interactions and characterizes them as specific interactions between the amphipathic faces of the MSPs and the solvent-membrane interfaces of the MPs. Our study provides structural and biophysical insights into how nanodiscs contribute to MP structural behavior and stability.

## RESULTS

### Membrane proteins have a spatial preference toward the edge of the nanodisc shells

López et al. used molecular dynamics simulation to investigate the behavior of the BAX protein incorporated into a nanodisc system <u>(López et al. 2019)</u>. Their analysis showed that the MP moved toward the edge of the nanodisc, where it was stabilized. Other studies showed this movement is entropy-driven <u>(Orioli et al. 2021)</u>. To further investigate the general spatial preference of MPs within the nanodisc shells, we revisited nine datasets of various MPs reconstituted in nanodiscs from EMPIAR (Table S1). We characterized the heterogeneity of the nanodisc shells using a two-step approach (Figure 1A). First, we generated consensus maps by aligning the MPs to establish a common reference frame using single particle analysis. Second, we classified the heterogeneity exhibited by the MSP relative to the MPs by focusing on the nanodisc shells. We observed an improvement in the intensity of the partial belt-like densities surrounding the MPs, which could potentially be attributed to the MSP. The analysis for all MPs displayed notable changes in the shape of the nanodiscs. Specifically, the nanodisc shells adopted an elliptical shape with varying degrees of ellipticity. Although the shape was altered, the overall volume of the nanodisc remained constant (Figure 1B). These findings collectively suggest that the MPs within the nanodiscs exhibit relative dynamics, which is reflected in their spatial distribution and the shape of the MSP.

**Figure 1.**
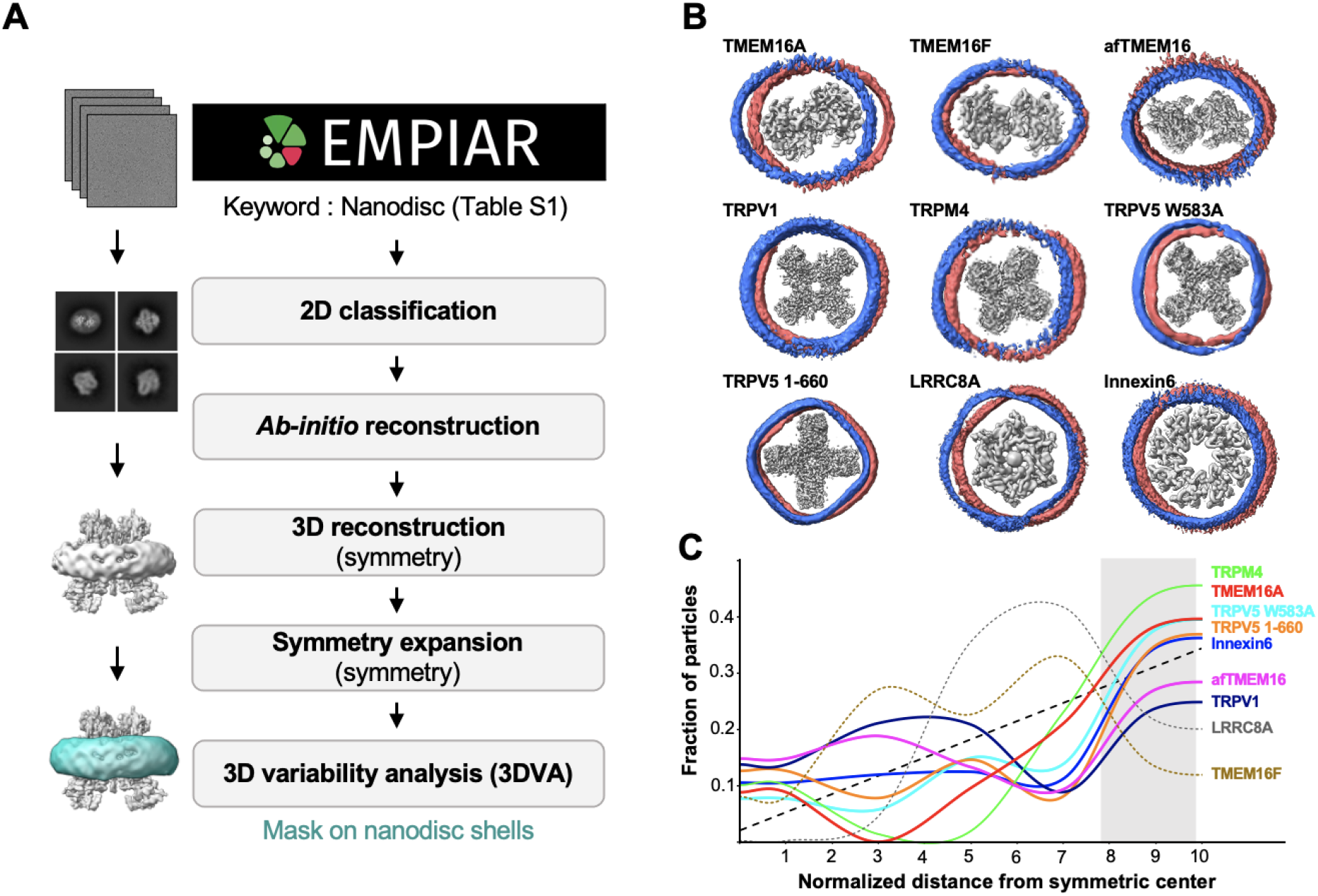
Analysis of the heterogeneous structure of the nanodisc. **(A)** Data processing workflow using CryoSPARC for 3DVA analysis. Among the membrane proteins (MPs) used in the analysis, TMEM16A is shown as a representative protein. **(B)** Results of 3DVA components show the subspace movement direction for the nine MPs used in the analysis. The relative position of the MSP to the MP in the first and last frames of the 3DVA are overlapped in blue and red. **(C)** The plot of the fraction of MP particles in each dataset that is a normalized distance away from the symmetric center of the nanodisc. The black dotted line represents the fraction of particles that would exist in each part of the nanodisc if the particles were to occupy each space homogeneously. The color-coded plot for nine MPs shows that for some, the fractions of particles farthest away from the center (gray) are larger than expected for the homogeneous distribution of particles (ex. TRPM4, TMEM16A, TRPV5, Innexin6). For such proteins, particles show unusually high occupancy at the edge of the nanodisc where the MSP may stabilize the MP.

To delineate the relative location of the MPs within the nanodiscs, we focused on discerning a subset of states that manifested linear motion of the MPs from one end of the MSP to the other while maintaining the overall nanodisc shape (Figure 1B, S1A). We scrutinized the particle population distribution using a quantitative model (Figure 1B, C, Figure S1C). By contrasting a control scenario positing homogeneous particle distribution within the nanodisc shells, our analysis revealed a pronounced enrichment of MPs at the periphery of the nanodisc shells (Figure 1C). Although the prevailing concept suggests that the MPs are predominantly localized in the central region of the nanodisc surrounded by the lipid molecules, this depiction may inaccurately imply that the protein is passively floating in the lipid environment. On the contrary, our collective observations revealed a more dynamic behavior, supporting previous observations of an entropic-driven migration of MP toward the nanodisc edge <u>(López et al. 2019; Orioli et al. 2021)</u>.

### Characterization of MP-MSP interactions

A significant portion of the acquired maps demonstrated ambiguous characteristics within the nanodisc shells, which can be ascribed to the inherent flexibility and dynamics of the MSP with the protein under investigation (Figure 2A). By applying a low-pass filter, we observed distinct belt-like density features predominantly located close to the membrane-solvent interface on the transmembrane helices of membrane proteins (MPs) (Figure 2A). These observations suggest that there are interactions between the MSP and MPs where the MSP may create local energy minima through their direct interactions. To characterize the interactions between the MPs and MSP, we analyzed the nanodisc-reconstituted maps from the EM database (Figure S2). Despite the limited resolution in the nanodisc shells that prevented the elucidation of the MSP-MP interactions in most cases, a notable exception was observed for the cytochrome bd-I oxidase protein map (EMD-4908) <u>(Safarian et al. 2019)</u>. This map exhibited a high-resolution density, resembling a segment of a single strand of MSP in direct contact with the cytochrome bd-I oxidase (Figure 2A). The fragment of the MSP map revealed a distinct density pattern consisting of eight α-helical turns, corresponding to 29 amino acids. Bulky side chain densities were observed at specific positions (n+3, n+10, and n+18, where n is the residue at the N-terminus) within the 29-residue segment. By excluding small residues (e.g. alanine, glycine, serine, or proline) at these positions, seven candidate sequences among MSP1D1 were selected and subsequently refined into the density (Figure 2B(i)). To find the best matching candidate, we employed Q-score analysis, which provides a quantitative assessment of the resolvability score for each atom and backbone (Pintilie et al. 2020). Our analysis revealed that the sequence 37-65 (QEFWDNLEKETEGLRQEMSKDLEEVKAKV) exhibited a significantly higher residue score compared to the other candidates (Figure 2B(ii)). This sequence had an optimal fit within the segmented MSP density (Figure 2B(iii)). Therefore, the observed fragment density corresponded to a specific region of MSP1D1, and this defined feature implied the presence of a constrained interaction between MSP1D1 and cytochrome bd-I oxidase.

**Figure 2.**
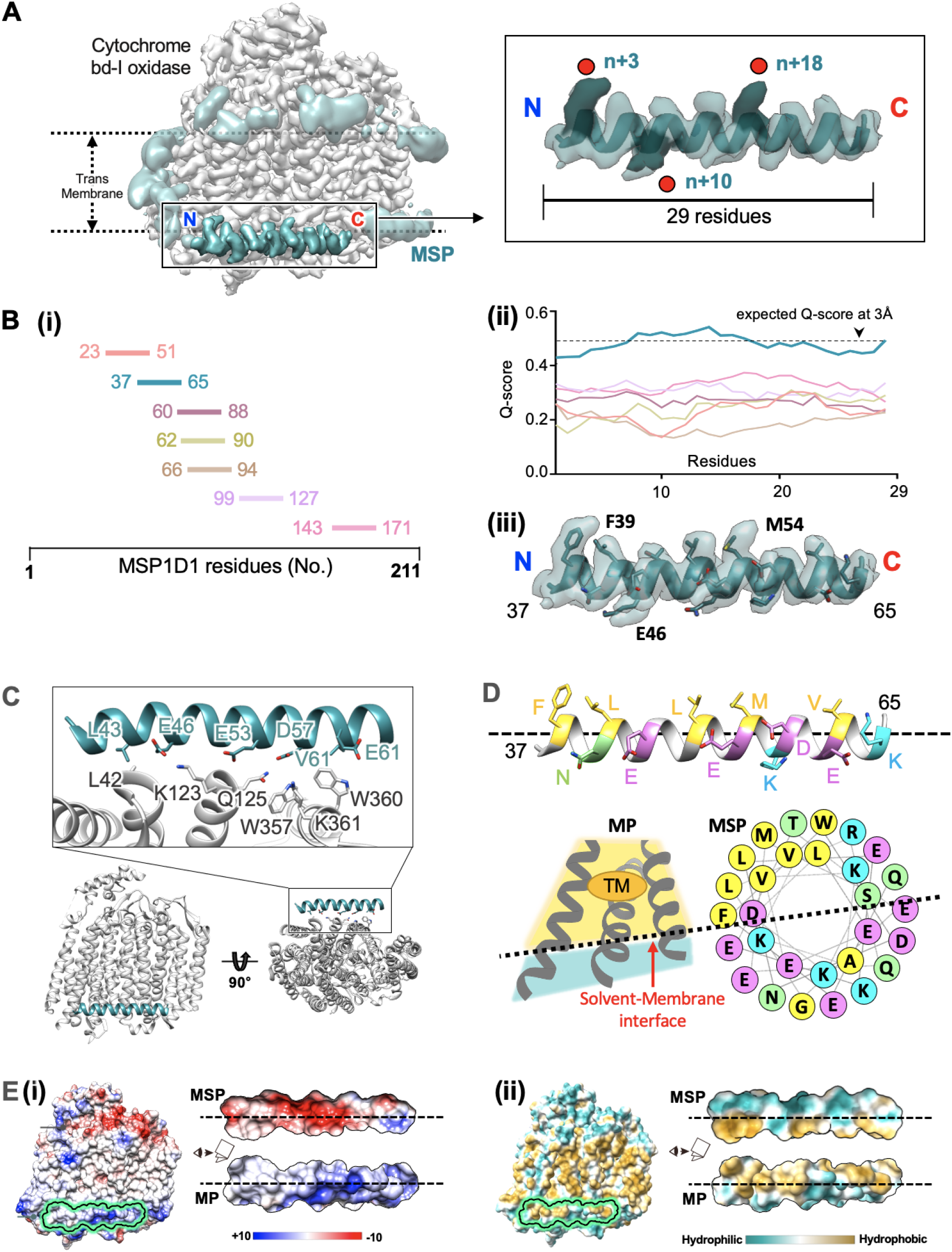
Characterization of MP-MSP interactions. **(A)** (*left*) Processed cryo-EM density map of cytochrome bd-I oxidase (gray) and MSP (cyan). The partial high-resolution MSP map on the edge of the transmembrane region of the MP is highlighted in the rectangular box. (*right*) A close-up of the high-resolution MSP region. Based on the number of helical turns, 29 residues are estimated to fit into the density. The bulky side chain densities of the n+3, n+10, and n+18 positions are highlighted. **(B)** (i) The seven 29-residue candidate sequences within MSP1D1 for best fit into the high-resolution MSP density map are color-coded. (ii) The Q-score for each candidate sequence is shown. The MSP sequence 37-65 (cyan) displayed the highest Q-score and was designated the best candidate for the high-resolution MSP density. (iii) The MSP sequence 37-65 best fits into the high-resolution MSP density. **(C)** Site of interaction between cytochrome bd-I oxidase (gray) and the high-resolution MSP (cyan) models. Residues potentially relevant to the interaction are highlighted in both the MSP and MP. **(D)** The helical wheel representation of sequence 37-65 shows the face of the MSP interacting with the cytochrome bd-I oxidase. Residues are color-coded based on their hydrophobicity, hydrophilicity, and charge. The MSP interacts with the MP at the solvent-membrane interface at the edge of the transmembrane region. **(E)** (i) (*left*) Electrostatic potential map of MP with the site of contact with the high-resolution MSP map highlighted. (*right*) The negatively charged residues on the contact face of the MSP can interact with the positively charged residues on the cytochrome bd-I oxidase. (ii) (*left*) Hydrophobic surface of the MP with the site of contact with the high-resolution MSP map highlighted. (*right*) A combination of the hydrophobic and hydrophilic regions in both MSP and the MP strengthen their interaction.

The MSP1D1 fragment was positioned perpendicularly to the luminal side of the membrane-solvent interface, and the helical pitches of the MSP matched well with the groove of the transmembrane helices CydX, CydA-H3, CydA-H4, and CydB-H9 of cytochrome bd-I oxidase. A meticulous examination of the interface revealed the presence of a combination of polar and hydrophobic residues (Figure 2C), resulting in a pronounced degree of polar and hydrophobic complementarity within the specific region. Notably, the MSP exhibited negatively charged residues, specifically E46, E53, D57, and E61, which emerged as potential candidates for interactions with the positively charged residues K123 and K361 within the MP. We also observed that the hydrophobic side of the MSP was oriented toward the hydrophobic transmembrane region of the MP (Figure 2E(ii)). Using PDBePISA, we also determined the thermodynamic binding energy, which was calculated as a negative free energy change of approximately 10 kcal/mol for this interaction. This energy change included the combinatorial contributions from the polar interactions (approximately -1.47 kcal/mol) and large solvation energy (-8.0 kcal/mol).

When considering the findings as depicted in a helical wheel representation (Figure 2D), the interactions between MSP and MPs are characterized by highly favorable binding between the amphipathic faces of the MSP and the solvent-membrane interface of the MPs. While this interaction could exhibit specificity (e.g. as observed with cytochrome), it might also display dynamic features with weak and heterogeneous constraints in most MPs (Figure S2).

### Nanodiscs contribute to membrane protein stability through MP-MSP interactions

We determined the influence of the close interactions of MSPs on the behavior and stability of the membrane proteins (MPs). For this purpose, we selected the vacuolar-type ATPase (V-ATPase) V_0_ complex from *S. cerevisiae* because its structure has been resolved under different conditions, including detergent (V_0_Det, 3.2 Å), amphipol (V_0_Amp, 3.9 Å), and nanodisc (V_0_ND, 2.7 Å) <u>(Pintilie et al. 2020; Vasanthakumar et al. 2019)</u>; <u>(Mazhab-Jafari et al. 2016)</u>; <u>(Roh et al. 2020)</u>. By comparing the features among the different maps, we observed a significant improvement in global resolution in the V_0_ND reconstruction compared to the other conditions (Figure 3A). Notably, while V_0_ND exhibited a uniform signal intensity throughout the entire map, the maps for V_0_Det and V_0_Amp displayed more varied signal intensities and higher B-factors, particularly on subunits c(3)-c(8) and d. Considering a similar number of particles were used for the final reconstruction, this enhanced global and local resolution in the V_0_ND reconstruction suggested that the particles reconstituted in the nanodiscs might possess greater structural stability.

**Figure 3.**
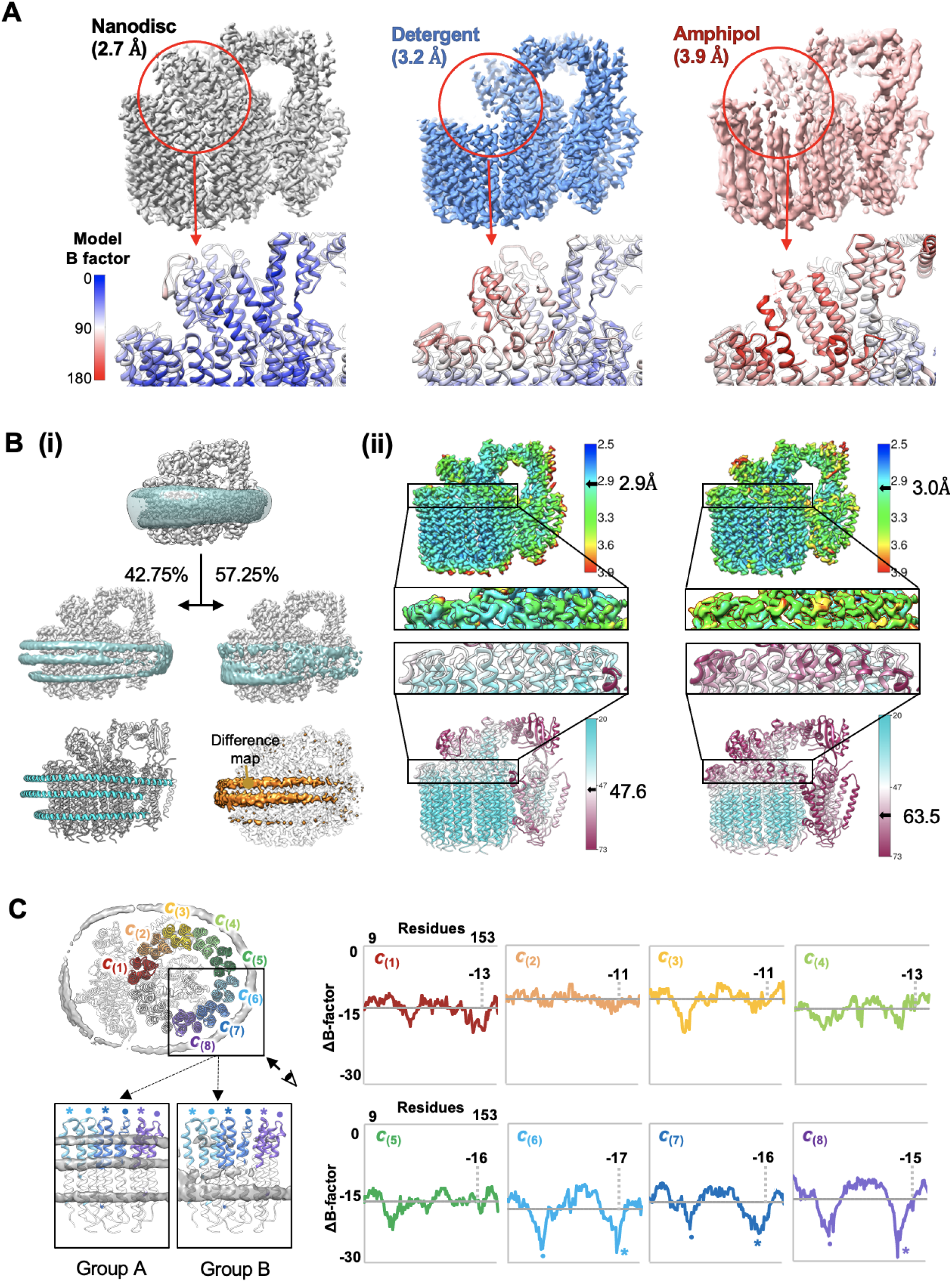
Nanodisc contributes to membrane protein stability. **(A)** Resolution and B-factor value comparisons among known cryo-EM structures of V-ATPase V_0_ complex in the nanodisc (*left*), detergent (*middle*), and amphipol (*right*). Specific regions in the structure with notable local resolution and B-factor differences are highlighted. **(B)** (i) Processed EM density map of the V_0_ complex in the nanodisc (*top*) was classified into two groups of particles for focused refinement. Group A (42.75% of total particles) contains particles with well-defined MSP features, while group B (57.25% of total particles) contains those with obscure or poorly-defined MSP features (*middle*). The three distinct MSP strands in the nanodisc surrounding the V_0_ complex are shown as a model (*bottom left*). The difference in the EM map between groups A and B (*bottom right*) isolates the MSP density responsible for improving the local resolution and B-factor in group A relative to group B. (ii) Resolution and B-factor value comparisons between group A (*left*) and B (*right*). The upper c-ring region with large deviations in local resolution and B-factor is highlighted. **(C)** (*left*) Top view of the V_0_ complex in the nanodisc with c-ring subunits color-coded. In the c(6), c(7), and c(8) subunits, the MSP density differences between groups A and B were the most conspicuous. (*right*) Plot of ΔB-factor values for each residue in the c-ring subunits is shown. ΔB-factor values were calculated by subtracting the B-factor values of group B from those of group A. The largest deviations were observed in the c(6), c(7), and c(8) subunits; the residues with the most noticeable differences are highlighted with dots and stars.

To gain further insights into the relationship between localized stability and the presence of the nanodiscs, we performed a comprehensive analysis of the V_0_ND particle images. By reconstructing the consensus map and applying a focused classification on the nanodisc shells, we separated the V_0_ particles and subsequently grouped them based on the density of the MSP features. Group A (43% of the total particles) exhibited three distinct and well-defined belt-like MSP densities, whereas group B (57% of the particles) displayed partial and weak MSP density (Figure 3B). The difference map between these groups clearly demonstrated the disparity in the upper two MSP densities, confirming the effectiveness of particle classification in detecting the presence of MSPs. Within group A, we observed that the top and bottom MSPs were closely positioned within about 10 Å of the V_0_’s c-ring, leaving limited space for lipid molecules. This observation suggested the potential for a direct interaction between the c-ring and the MSPs.

The 3D reconstruction of both groups exhibited distinct and well-defined MP features, with overall average resolutions of approximately 2.9 Å for group A and 3.0 Å for group B (Figure 3B). However, a closer analysis of the local areas between the two groups revealed a significantly higher resolution and a more stable average B-factor in group A, specifically in the cytosolic side of the c-ring region (Figure 3B). Interestingly, this region aligned with the cytosolic region of the c-ring, where a noticeable discrepancy in resolution was observed between V_0_ND and V_0_Det (Figure 3A). This consistent finding suggested that defined MSP interactions play a role in stabilizing the MP.

We also analyzed the per-residue B-factor changes (ΔB-factor values) between groups A and B to assess the degree of stabilization resulting from the MSP interactions. For each residue, the ΔB-factor value was calculated by subtracting the B-factor value of group B from that of group A. We focused on eight identical c-subunits (c(1) through c(8)) that exhibited varying exposure within the nanodisc shells (Figure 3C(i)). Notably, subunit c(1) interacted with subunit a, whereas c(2) was partially exposed to the phospholipid and had limited contact with the MSP. In contrast, c(3) through c(8) were exposed to the MSPs. When comparing chains c(1) and c(2), where the MP does not directly interact with the MSPs, the alterations in the B-factors had no significant distinctions (Figure 3C(ii)). Conversely, subunits c(3) through c(8) displayed notable negative ΔB-factor values on average. Subunits c(6), c(7), and c(8) exhibited especially large deviations in their B-factor values (Figure 3C(ii)). Taking a closer look at the structure, we also observed the largest difference in MSP features between groups A and B in the c(6), c(7), and c(8) subunits. In group A, we observed three distinct MSP strands surrounding the residues in these subunits, whereas we observed a distinct MSP feature in group B only in the bottom strand (Figure 3C(i)). Correspondingly, the residues in the c(6), c(7), and c(8) subunits with the largest ΔB-factor values were in the cytosolic region that interacted with the top strand of the MSP (Figure 3C(ii), dot and star). Taken together, these data demonstrated that the stability of the MSP surrounding the MP conferred stable direct interactions with the MP itself in those regions to lower their B-factor values and enhance protein stability. These findings suggest that direct interaction with the MSP could enhance protein stability.

### Multivalent MSPs contribute to membrane protein stabilization

Since MSP is derived from apolipoprotein A1, which consists of two aligned strands, it has been commonly assumed that MSPs in nanodiscs adopt a configuration with two strands. However, we observed three MSP belts within the V-ATPase V_0_ complex. The top and bottom MSPs were positioned near the interfaces between the membrane and solvent in V_0_. The middle MSP strand was located at a greater distance from the V_0_ but at an approximate distance of 15 Å from the top MSP, corresponding to the typical spacing between helices in apolipoproteins (Figure 4A). Analysis of the database also showed three MSPs in glucosyltransferase Alg6 (PDB 6SNI, EMD-10258) and arabinofuranosyltransferase D (AftD) (PDB 6W98, EMD-21580) (Figure S2). Given that the transmembrane domains of membrane proteins (MPs) typically exceed a length of 30 Å, our findings indicated a reorganization of the interactions from MSP-MSP into MSP-MP. Consequently, the presence of an additional MSP might serve to shield the lipid environment surrounding the MPs (Figure 4B). This observation implied that the MSPs engaged in a dynamic interplay involving both traditional and complementary interactions with neighboring MSPs and direct interactions with MPs. Such a combination of interactions could preserve nanodisc morphology while simultaneously accommodating lipids and stabilizing the MPs.

**Figure 4.**
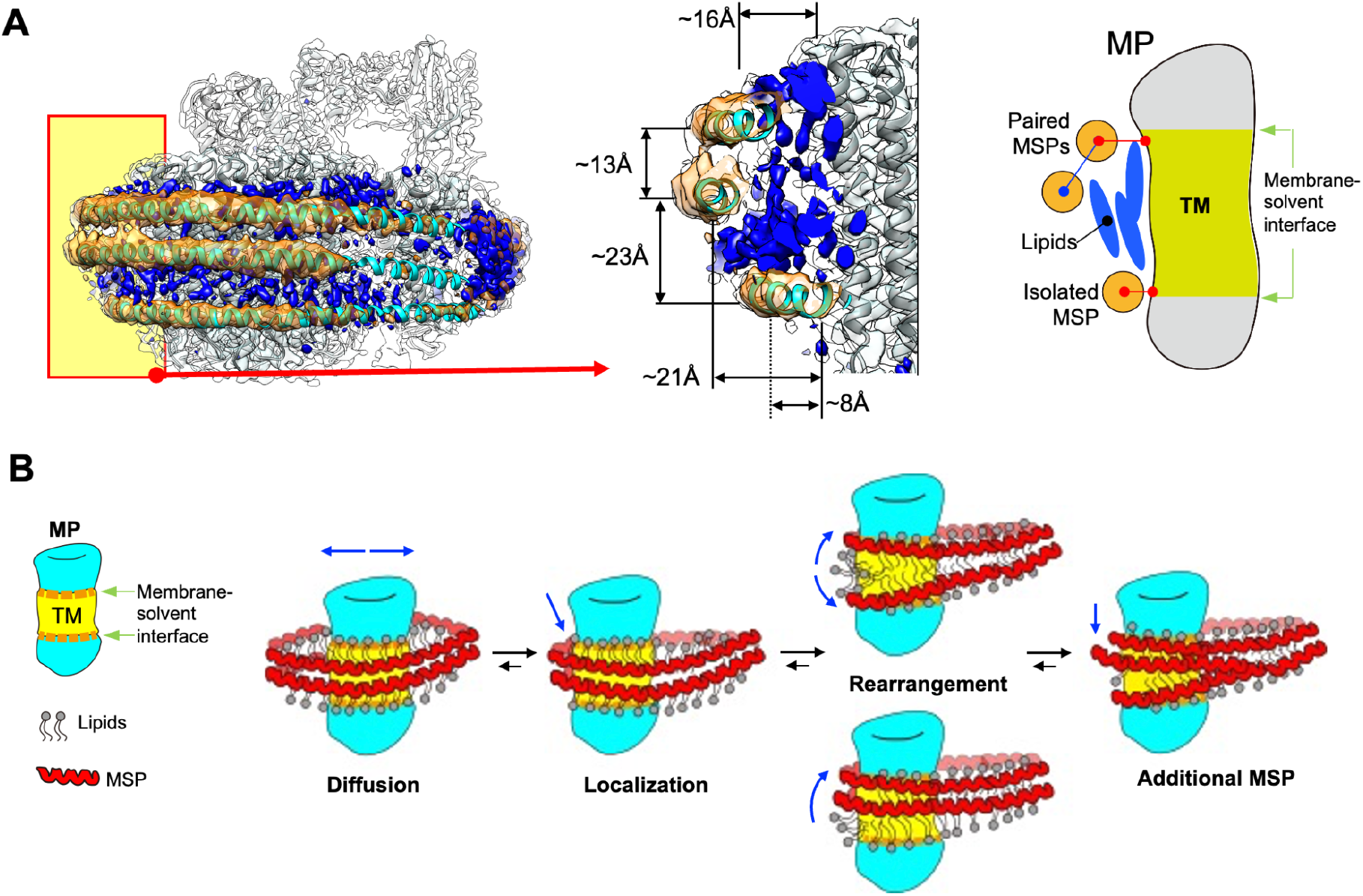
Multivalent MSP binding to membrane proteins and stabilizing mechanism. **(A)** (*left*) V-ATPase V_0_ complex model (gray) and density map (white), including the MSP model (cyan) and map (orange). The density maps corresponding to the lipid molecules are colored in blue. The site of the MP-MSP interactions is highlighted with a yellow box for a zoomed-in view (*middle*). The distances between the top, middle, and bottom MSP strands and membrane protein (MP) were approximately 16 Å, 21 Å, and 8 Å, respectively. The distance between the top and middle MSP strands was about 13 Å, while that between the middle and bottom strands was approximately 23 Å. The lipid molecules could occupy the space between the top and middle MSP strands and the V_0_ complex but not the space between the bottom MSP strand and the V_0_. (*right*) A schematic view of the MP-MSP interactions. The top and bottom MSP strands surround the MP at the membrane-solvent interface. The bottom strand forms a direct and stable interaction with the MP through a lipid-free local environment (red line). The top strand interacts with both the middle strand of the MSP (blue line) and the MP (red line). **(B)** The schematic illustration depicts the nanodisc stabilization process after initial assembly. The MSP bilayer (red) envelops the MP (cyan) and phospholipid (PL) (gray), supported by salt-bridge interactions between the MSP strands. The MP undergoes diffusive movement within the nanodisc, and the MP and MSP can make contact. The MSP-MSP salt-bridge interaction is then converted to MP-MSP interactions. The overall nanodisc adopts an elliptical shape during this process. If the MP has a TM length above a certain threshold, the two strands of the MSP come far enough apart to allow the presence of a third MSP strand, further stabilizing the MP through MP-MSP contacts.

## DISCUSSION

The use of nanodiscs for the reconstitution of membrane proteins (MPs) has gained considerable popularity as a purification method for biophysical and biochemical studies. In our experimental analysis, we observed that MPs exhibited a spatial preference near the edges of the nanodisc shells, allowing them to establish contact with the MSPs. These interactions are dynamic, occurring between the amphipathic surfaces of the MSPs and the interface between the solvent and the membrane of the MPs. This preference for the nanodisc periphery can perhaps be attributed to the relatively thinner lipid bilayer in this region compared to the center of the nanodisc <u>(Maingi and Rothemund 2021)</u>.

Compared to purification in detergents, nanodisc-reconstituted MP structures have been associated with higher resolution <u>(Autzen et al. 2019)</u>. Our study provides insights into the stabilizing effect of nanodisc structure on MPs. While the scientific community generally considers nanodiscs to provide a more native-like environment for proteins, we observed some protein-protein interactions when MPs were reconstituted in nanodiscs. In fact, the presence of such interactions has been observed by Kern et al. in their cryo-EM structure of the nanodisc-reconstituted SARS-CoV-2 ORF3a protein. They also observed partially well-resolved MSP1E3D1 density where MSP interacts with the MP <u>(Kern et al. 2021)</u>. Recently, Dalal et al. also observed that the use of varying MSP sizes influenced the conformational states of the reconstituted protein. The smaller MSP tends to bind the protein more tightly, altering the properties of the lipid bilayer or directly interacting with the protein of interest, while larger MSP tends to provide a more structurally supportive role by providing a native environment <u>(Dalal et al. 2022)</u>. Nonetheless, the implications of these MP-MSP interactions on the physiological function of the purified proteins remain elusive and require extensive investigation for validation. However, these characterized interactions from previous literature as well our findings in this study may assist researchers in selecting appropriate nanodiscs for their experiments.

From our analyses, we propose the characteristics of MPs reconstituted in nanodiscs (Figure 4B). During the initial reconstitution of MPs in nanodiscs, the hydrophobic regions of the MPs are reconstituted by the phospholipid bilayer, which is stabilized by the surrounding MSP proteins. The MSP strands are initially stacked to each other and stabilized by a complementary charge interaction <u>(Klon et al. 2002)</u>. MPs in the nanodisc undergo lateral diffusion and are entropically stabilized at the nanodisc edge <u>(Orioli et al. 2021)</u>. MPs establish contacts with the proximal MSPs, converting MSP-MSP strand interactions into MP-MSP interactions. During this reorganization, each MSP shifts to the membrane-solvent interface of the MPs. Due to the amphipathic nature of MSPs, both hydrophobic and charge-charge interactions form between the MPs and MSPs, generating a stabilized MP-embedded nanodisc. As the two strands of MSP are rearranged to the membrane-solvent interfaces on the MPs, the hydrophobic TM region of the MPs can be exposed to an aqueous solution, which may destabilize the protein. To accommodate this effect, an additional MSP strand surrounds the exposed hydrophobic region for stabilization. Depending on the TM length of the MP, the MPs may adopt three strands rather than two MSPs for stabilization.

## ACKNOWLEDGMENTS

This work was supported by the Korean National Research Foundation (2019R1C1C1004598, 2020R1A5A1018081, 2021M3A9I4021220, and 2019M3E5D6063871) and the Suh Kyungbae Foundation (SUHF) to S.-H.R. We thank members of Center for Macromolecular and Cell Imaging (CMCI) for their discussions and suggestions.

## AUTHOR CONTRIBUTIONS

Conceptualization, S.-H.R.; investigation, S.-J.K., Y.H.K.; writing, S.-J.K., Y.H.K., and S.-H.R.; Funding Acquisition, S.-H.R.

## AUTHOR CONTACTS

So-Jung Kim - Young Hoon Koh - Soung-Hun Roh -

## MATERIALS AND METHODS

### Cryo-EM image processing

All raw data for the analysis were obtained from EMPIAR (https://www.ebi.ac.uk/empiar/). Processing was done using CryoSPARC <u>(Punjani et al. 2017)</u>. The input values from each dataset are listed in **Table S1**. We used MotionCor2 for motion correction of images and CTFFIND4 for CTF estimation <u>(Rohou and Grigorieff 2015)</u>. Initial 2D templates were generated using a blob picker. Additional rounds of particle picking were carried out via template-based picking. The initial 3D model with C1 symmetry was built based on the reference 2D classes. The initial map was refined through iterative non-uniform (NU) and heterogeneous refinement processes. Once the desired resolution was reached, global and local CTF refinement followed by additional rounds of NU refinement were performed. For heterogeneity analysis of the nanodiscs, we first isolated the nanodisc density from the final map using Chimera’s segment map and map eraser features <u>(Pettersen et al. 2004)</u>. We then imposed a maximum value of the Gaussian volume filter to erase the map details. A mask of the nanodisc region was created from CryoSPARC with a dilation radius and soft padding width of 3 pixels. After final processing, we performed symmetry expansion on the particles. In conjunction with the nanodisc density map, output results were used as input for 3D Variability Analysis (3DVA) <u>(Punjani and Fleet 2021)</u>. A filter resolution of 7 was used to solve three modes of variability.

### Quantitative analysis of heterogeneity within a protein-embedded nanodisc

For quantitative analysis of heterogeneity in protein-embedded nanodiscs, each MP dataset was put into the pipeline for 3DVA (Figure 1A). Raw data downloaded from the EMPIAR database were subjected to 2D classification, which was used for *ab-initio* reconstruction. The initial 3D model was used as a model for 3D reconstruction, which was fed into the 3DVA for the symmetry expansion process. The 3DVA output cluster was divided into 20 frames. We quantified the location of the MP relative to the MSP and the varying changes in the shape of the nanodisc itself. Five different variables were considered for our study (Figure S1C). As a caveat, we assumed the nanodisc adopted an elliptical shape in its 2D projection. We considered only the 2D projection rather than the entire 3D structure because the shape of the nanodisc is not an ellipsoid in 3D but rather an elliptical cylinder. The variables considered for the nanodisc included: the radius of the long and short axes of the ellipse, denoted as “a” and “b,” respectively, and the focal point of the nanodisc “c.” The sign of “c” referred to the direction of elliptical deformation, whereas the magnitude of “c” described the degree of ellipticity of the nanodisc. We considered the variables describing the position of the MP within the nanodisc relative to the MSP as “θ” and “ΔR,” where “θ” was the angle formed between the long axis “a” and the line connecting the center of the nanodisc to the center of the MP, and “ΔR” was the distance between the center of the nanodisc to that of the MP.

We analyzed three orthogonal yet complete “modes” from the 3DVA. For each mode, the motion of the MP relative to the nanodisc was such that all but one variable was fixed. We normalized the maximum value at 10 for each case and binned by five intervals. The homogeneous distribution graph was calculated for each variable. For instance, for modes where ΔR was allowed to vary but other variables (e.g., θ and c) were fixed, the cross-section of the nanodisc formed either a fixed circle or an ellipse, depending on the given c value. For c = 0, any frames of the 3DVA with the same ΔR value were degenerate as they were indistinguishable from one another. In this case, the homogeneous distribution graph was calculated by adding the degeneracies together within the range of normalization, with nR referring to the normalized value of R (for 0 < nR < 2, 1/25 of total particles, for 2 < nR < 4, 3/25 of total particles, for 4 < nR < 6, 5/25 of total particles, for 6 < nR < 8, 7/25 of total particles, and for 8 < nR < 10, 9/25 total particles) (Figure S1C). When c was taken as the normalized variable (nc), even if the nR value was the same, each frame was uniquely defined for each “c” value. Thus, this case produced a homogeneously distributed graph (Figure S1C).

### Structure visualization

We displayed published structural maps of various MPs using the Coulombic surface coloring feature in Chimera to highlight the surface charges. We applied a 3 Å low-pass filter to the nanodisc density and the MP in CryoSPARC. In Chimera, we separated the density of the MSP from the filtered map through a segmentation process.

### Identification of the sequence fragment in MSP1D1

We isolated only the EM density corresponding to the MSP from a 3 Å low-pass filtered map of the cytochrome bd-I oxidase protein using the same process described above. The map was re-segmented to observe the side chain density map of the MSP. The candidate sequence came from the full sequences of MSP1D1 used in the nanodisc reconstitution of cytochrome bd-I oxidase (Figure 7A). The high-resolution MSP map was composed of an eight-turn α-helix. Based on 3.6 amino acids per turn of α-helix, we estimated the total length as 29 amino acids. To determine the candidate sequences within the MSP, we used the 3^rd^, 10^th^, and 18^th^ bulky amino acids from the MP contact side as a reference within the high-resolution MSP map (Figure 7B). For the fitting, the 3^rd^ residue in the chain was designated F, K, Y, or R as the bulkiest side chains, which left 30 candidate MSP sequences that might fit into the MSP map. The 10^th^ and 18^th^ residues also showed bulky side chain densities, allowing the elimination of residues A, P, G, and V occupying those sites. Through this process of elimination, we had seven final candidate sequences (Figure 7C). Initial model fitting for all seven candidate MSP sequences was done using ISOLDE, an MD simulation-based software <u>(Croll 2018)</u>. We used the default setting values for all parameters except for α-helix restrain. We improved the accuracy of the model fitting using real-space refinement software from Phenix <u>(Croll 2018; Adams et al. 2010)</u>. We calculated the Q-score from the fitted models of density resolvability for protein and quantitative evaluation <u>(Pintilie</u> <u>et al. 2020)</u>. In the 3.0 Å filtered map, the Q-score value for each candidate sequence was compared to the average Q-score value to determine the best candidate.

### Cytochrome BD-I oxidase and MSP binding energy calculation using PDBePISA

We conducted PDBePISA analysis to gain thermodynamic insights into the MP-MSP contact interface <u>(Krissinel and Henrick 2007; Krissinel 2010)</u>. The software identified three hydrogen bond interactions (K361:D57, Q125:E53, and Q125:D57) and one salt-bridge interaction (K361:D57). The free energy contribution from the hydrogen bond and electrostatic binding interactions was Δ^P^G = Δ^H^G + Δ^e^G = ((-0.44 x 3) + (-0.15)) kcal/mol = -1.47 kcal/mol. The software also calculated the solvation-free energy of Δ^S^G = -8.0 kcal/mol, indicating spontaneous embedding through hydrophobic interfaces. In total, the free energy of the MP-MSP interaction was estimated to be ΔG = Δ^P^G + Δ^S^G = -9.47 kcal/mol. Therefore, the interaction between cytochrome bd-I oxidase and the MSP was favorable and spontaneous. PDBePISA also provided Δ^S^G P-value of P = 0.346. The P-value was the probability of obtaining a solvation energy gain lower than the observed (-8.0 kcal/mol) if the interface atoms were picked randomly from the protein surface. It measures interface specificity. P = 0.346 indicated that the interface contained notable hydrophobicity, implying that the interface surface could be interaction-specific.

### V-ATPase V_0_ image analysis

The V-ATPase V0 image analysis was carried out with CryoSPARC, with the first 3D reconstruction of the V-ATPase V_0_ complex performed in RELION 3.0 <u>(Pintilie et al. 2020;</u> <u>Zivanov et al. 2018)</u> with a 2.81 Å resolution. The 3D classification included 476k processed particles, the map from CryoSPARC, and the mask of the nanodisc region of the V_0_ complex depicted using Chimera. A regularization parameter T of 4 was used with no image alignment. The resulting classes were categorized by whether or not the MSP strands were visible. The high-resolution MSP group had 203,419 particles (42.75% of total particles), whereas the low-resolution MSP group had 272,394 particles (57.25%). Using the NU refinement program from CryoSPARC, three sets of 3D reconstructions were performed with the total number of particles, the high-resolution MSP group, and the low-resolution MSP group. The input volume was the same as the map used in RELION. The resulting output volume and the mask of only the MP region were used as input for the local refinement process in RELION to improve local resolution in the MP region. The total particles, high-resolution, and low-resolution groups had final resolutions of 2.86 Å, 2.92 Å, and 2.99 Å, respectively.

### Quantitative analysis for structural stability

In CryoSPARC, we calculated the local resolution of the MSP using the output volume and mask derived from local refinement. In Chimera, MSP surface coloring was performed from the output volume of the local refinement process. The local resolution output volume was used as the input volume data value. The color key was set in five intervals (2.5, 2.9, 3.3, 3.6, and 3.9 Å). In Phenix, we performed real-space refinement with default settings using the V-ATPase V_0_ complex map and model (6M0R) as the input. From this refinement, we obtained the atomic displacement parameter value (B-factor) for each residue in the protein.

## Data sharing plans

Further information and requests for resources and reagents should be directed to and will be fulfilled by the Lead Contact, Soung-Hun Roh.

## LIST OF SUPPLEMENTAL INFORMATION

**Table S1.** Datasets utilized from EMPIAR/EMDB/PDB Database.

| Protein |  |  | EMPIAR/<br>EMDB/<br>PDB | Published<br>Resolution (Å) | Reprocessing<br>Resolution (Å) | MSP | Lipid | MP:MSP:Lipid | Reference |
| --- | --- | --- | --- | --- | --- | --- | --- | --- | --- |
| TRP family | TRPV1 |  | 10059/<br>8117/<br>5IRX | 2.95 | 2.79 | 2N2 | Soy PC | 1:1:150<br>~ 1:1.5:225 | (Gao et al. 2016) |
|  | TRPV5 | 1-660 | 10255/<br>0593/<br>6O1N | 2.9 | 2.88 | 2N2 | Soy PC | 1:2.5:100 | (Dang et al. 2019) |
|  |  | W583A | 10253/<br>0605/<br>6O1U | 2.85 | 2.85 | 2N2 | Soy PC | 1:1.5:150 | (Dang et al. 2019) |
|  | TRPM4 |  | 10127/<br>7133/<br>6BQV | 3.1 | 2.86 | 2N2 | Soy PC | 1:3:390 | (Autzen et al. 2018) |
| TMEM 16<br>family | TMEM16A |  | 10123/<br>7095/<br>6BGI | 3.8 | 3.48 | 2N2 | Soy PC | 1:4:100 | (Dang et al. 2017) |
|  | TMEM16F |  | 10280/<br>20246/<br>6P48 | 3.3 | 3.06 | 2N2 | Soy PC:<br>POPC:<br>POPE:POPS<br>= 6:3:3:1 | 1:4:100 | (Feng et al. 2019) |
|  | afTMEM16 |  | 10240/<br>8959/<br>6E1O | 3.59 | 3.83 | 1E3 | POPE:POPG<br>= 3:1 | 1:2.5:250 | (Falzone et al. 2019) |
| Anion channel | LRRC8A |  | 10258/<br>0562/<br>6NZW | 3.21 | 3.09 | 1E3D1 | POPC | 1:2.5:250 | (Falzone et al. 2019; Kern et al. 2019) |
| Hemichannel | Innexin6 |  | 10291/<br>9973/<br>6KFH | 3.6 | 3.37 | 2N2 | POPC | 1:0.5:30 | (Burendel et al. 2020) |
| Cytochrome bd-I oxidase |  |  | -/<br>4908/<br>6RKO | 2.68 | - | 1D1 | POPC | 1:10:400 | (Safarian et al. 2019) |
| V-ATPase V<sub>0</sub> complex |  |  | -/<br>30035/<br>6M0R | 2.7 | 2.82 | 1E3D1 | *E.coli*<br>total lipid<br>extract | 1:50:1250 | (Roh et al. 2020) |
| Alg6 |  |  | -/<br>10258/<br>6sni | 3.0 | - | 1D1 | DDM:CHS | 1:6:390 | (Bloch et al. 2020) |
| AftD |  |  | 10399/<br>21580/<br>6w98 | 2.9 | 2.99 | 1E3D1 | *E.coli*<br>total lipid<br>extract | 1:6:360 | (Tan et al. 2020) |

**Figure S1.**
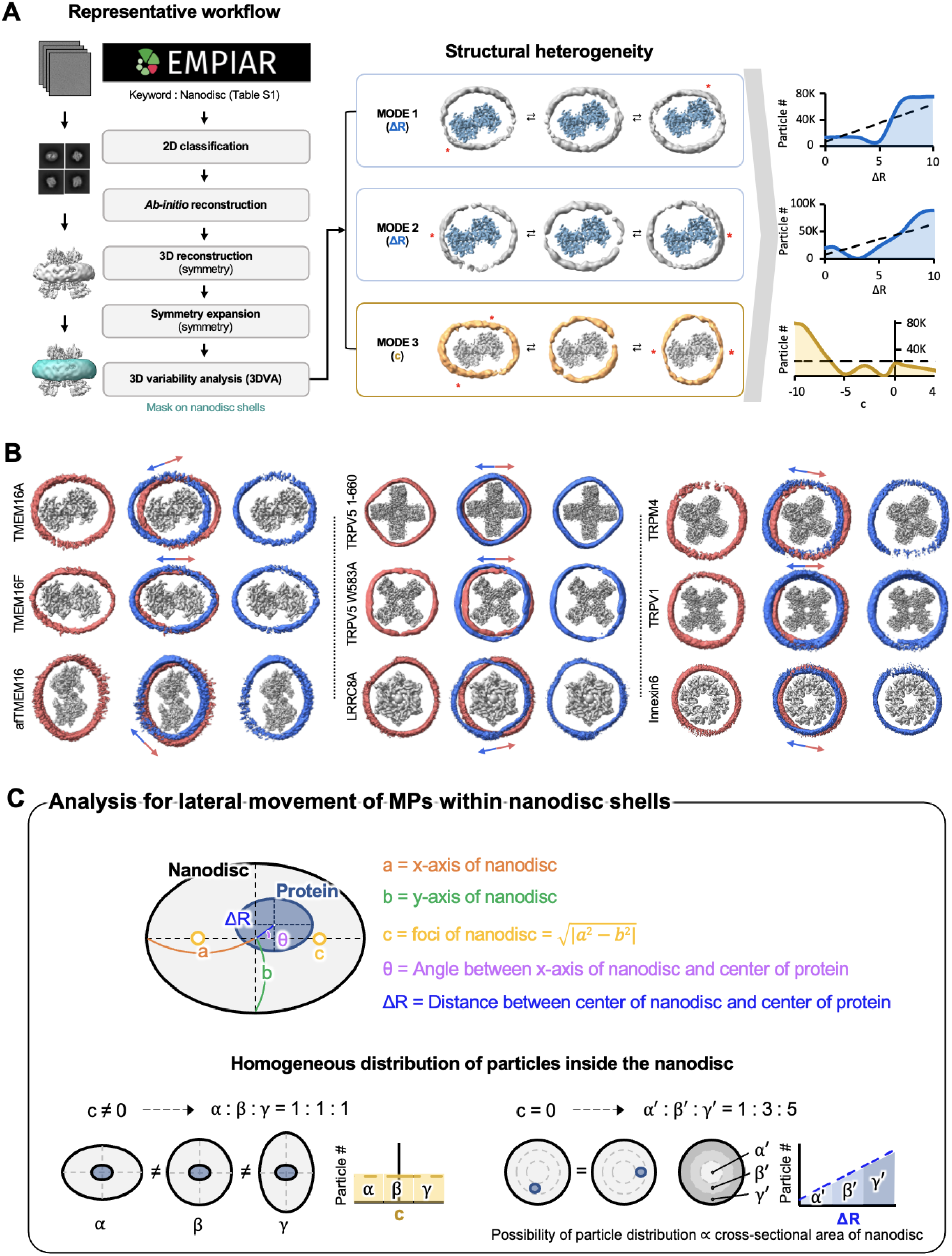
Schematic of the quantitative model used to analyze the distribution of membrane protein within the nanodisc. (A) (*left*) Representative workflow of 3D Variability Analysis (3DVA). Raw data obtained from EMPIAR was processed for 2D classification. The results were used for *ab-initio* reconstruction, which was used as a model for 3D reconstruction. Symmetry expansion was performed before feeding the structure in 3DVA. (*middle*) Three modes of structural heterogeneity were isolated from the 3DVA. In modes 1 and 2, the membrane protein (MP) moves linearly from one edge of the MSP to the other (ΔR). This movement is orthogonal for modes 1 and 2, and the shape of the nanodisc remains the same during the movement. In mode 3, the MP remains the same while the nanodisc deforms into an elliptical shape. The degree of deformation was estimated by the value c, the focal point. TMEM16A is shown as an example. (*right*) For each 3DVA mode, particle distribution was calculated and plotted as a function of either ΔR or c. The dotted lines represent the particle distribution for homogeneously distributed particles for each ΔR or c. The solid lines show the result for TMEM16A. In modes 1 and 2, the particles are distributed more at the edge and the center of the nanodisc and less in between than expected. In mode 3, the particles are highly populated in a state where the nanodisc is deformed heavily such that the MSP can come into contact with the MP itself. (B) One mode subspace movement is shown for the nine MPs used in the analysis. The relative positions of the MSP to the MP in the first and last frames of the 3DVA are shown in blue and red. The arrows show the directions of the movement. (C) In the analysis of the movement of MPs within the nanodisc, we defined necessary variables (a, b, c, θ, ΔR’) for our MP-embedded nanodisc model (see Materials and Methods). As a control, homogeneous distributions of the particles inside the nanodisc were considered. For c ≠ 0, each value of c is equally likely, and thus we should obtain a horizontally flat line in the number of particles vs. c value graph. For c = 0, particles with a particular value of ΔR are degenerate. The value of this degeneracy increases as R increases. Therefore, we expect to observe an R-dependent linear increase in particle distribution in the number of particles vs. ΔR graph.

**Figure S2.**
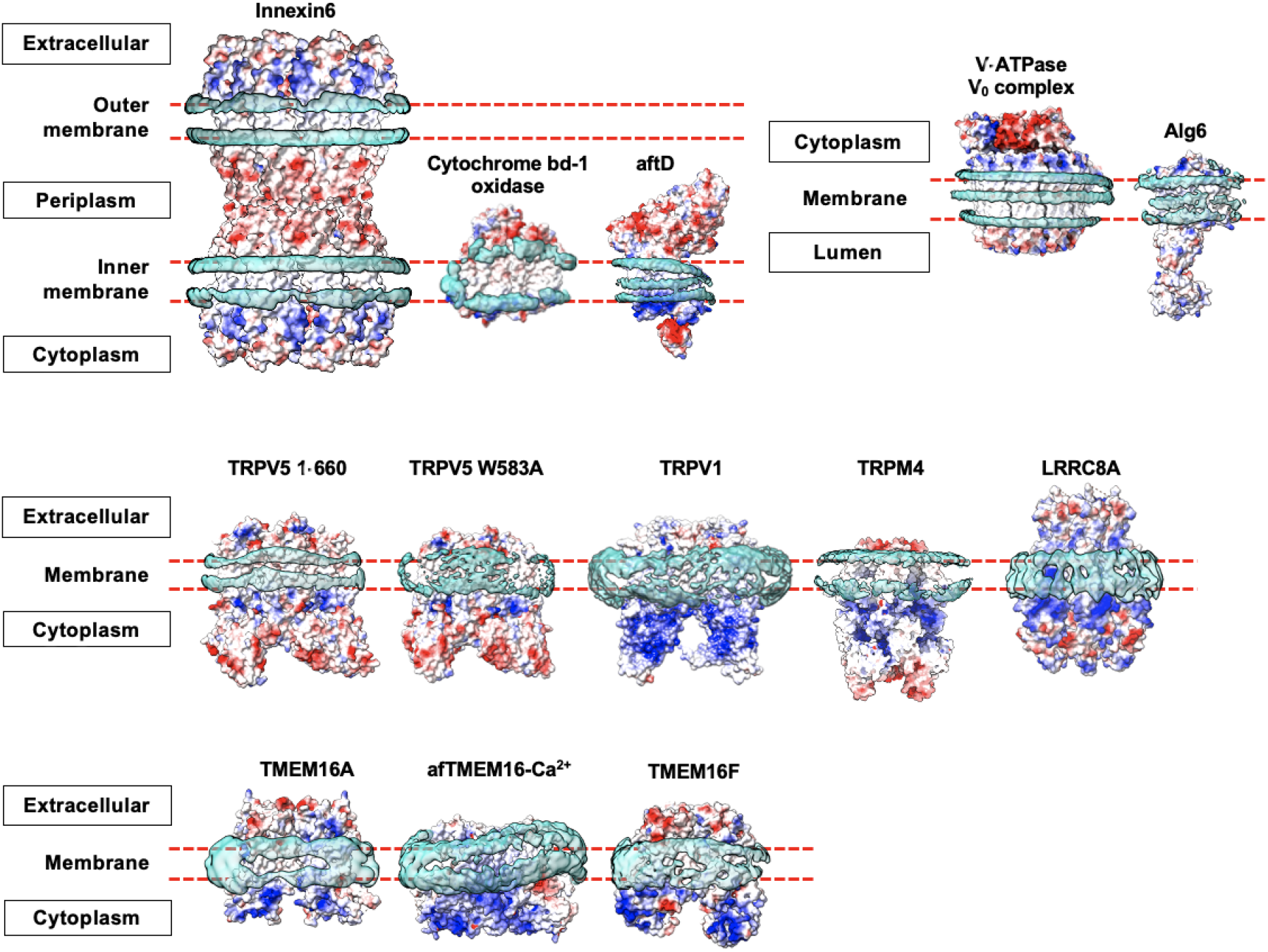
Nanodisc shells at low-threshold display. Membrane proteins (MPs) used in the analysis are presented with their surface charges colored. Each protein is classified according to its respective location in the cell. Low-threshold densities attributed to the MSPs are shown in sky blue.

**Figure S3.**
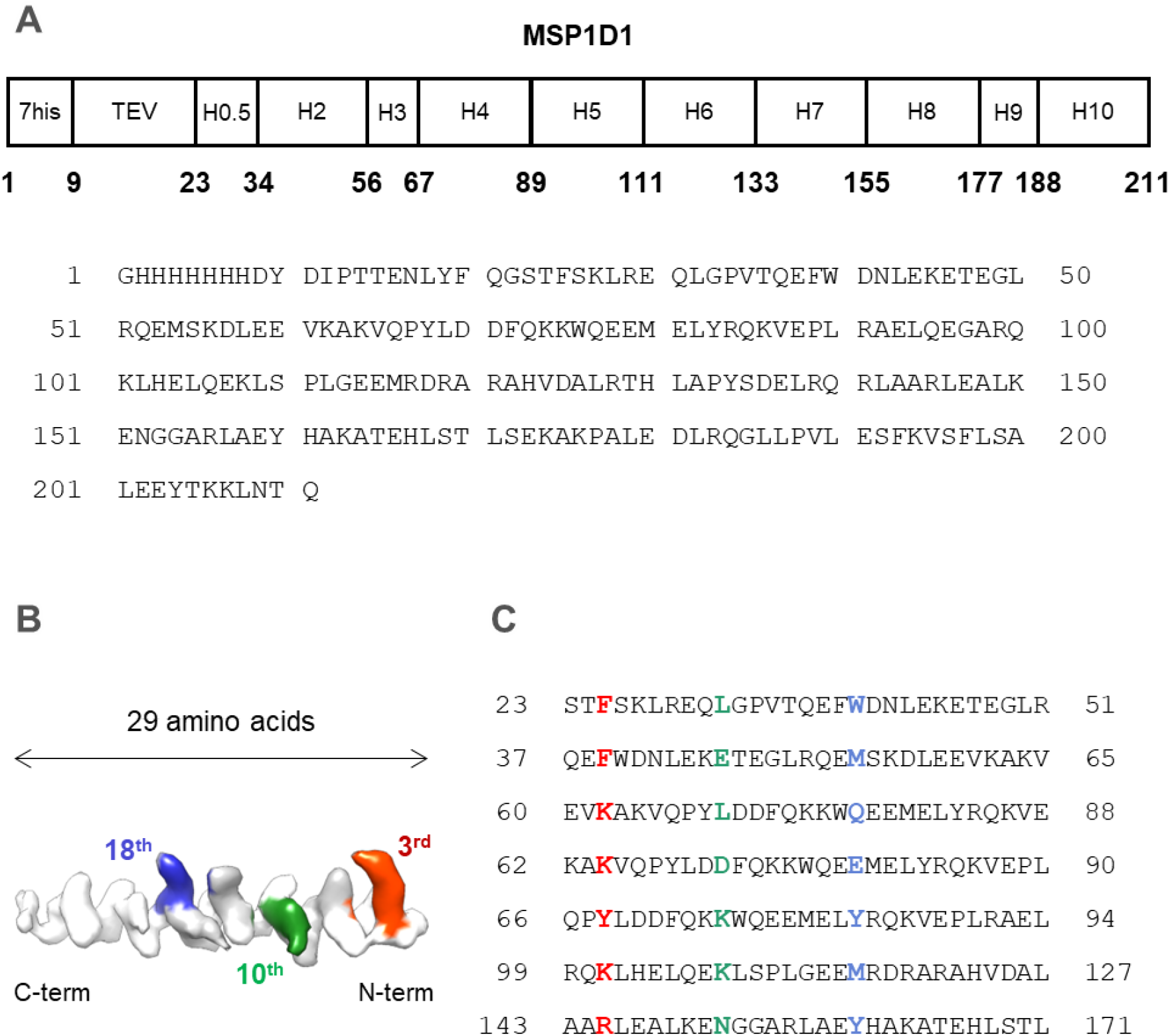
The amino acid sequence of MSP1D1 and candidates used for model fitting. (A) MSP1D1 construct map and full amino acid sequences. (B) Partial high-resolution MSP map of cytochrome bd-I oxidase. The MP-MSP contact side is shown, with the C-terminal and N terminal end on the left and the right, respectively. Colored residues represent bulky side chain density maps used as a reference for selecting candidate sequences for model fitting. (C) Seven candidate sequences were tried for model fitting into the MSP high-resolution map.

**Figure S4.**
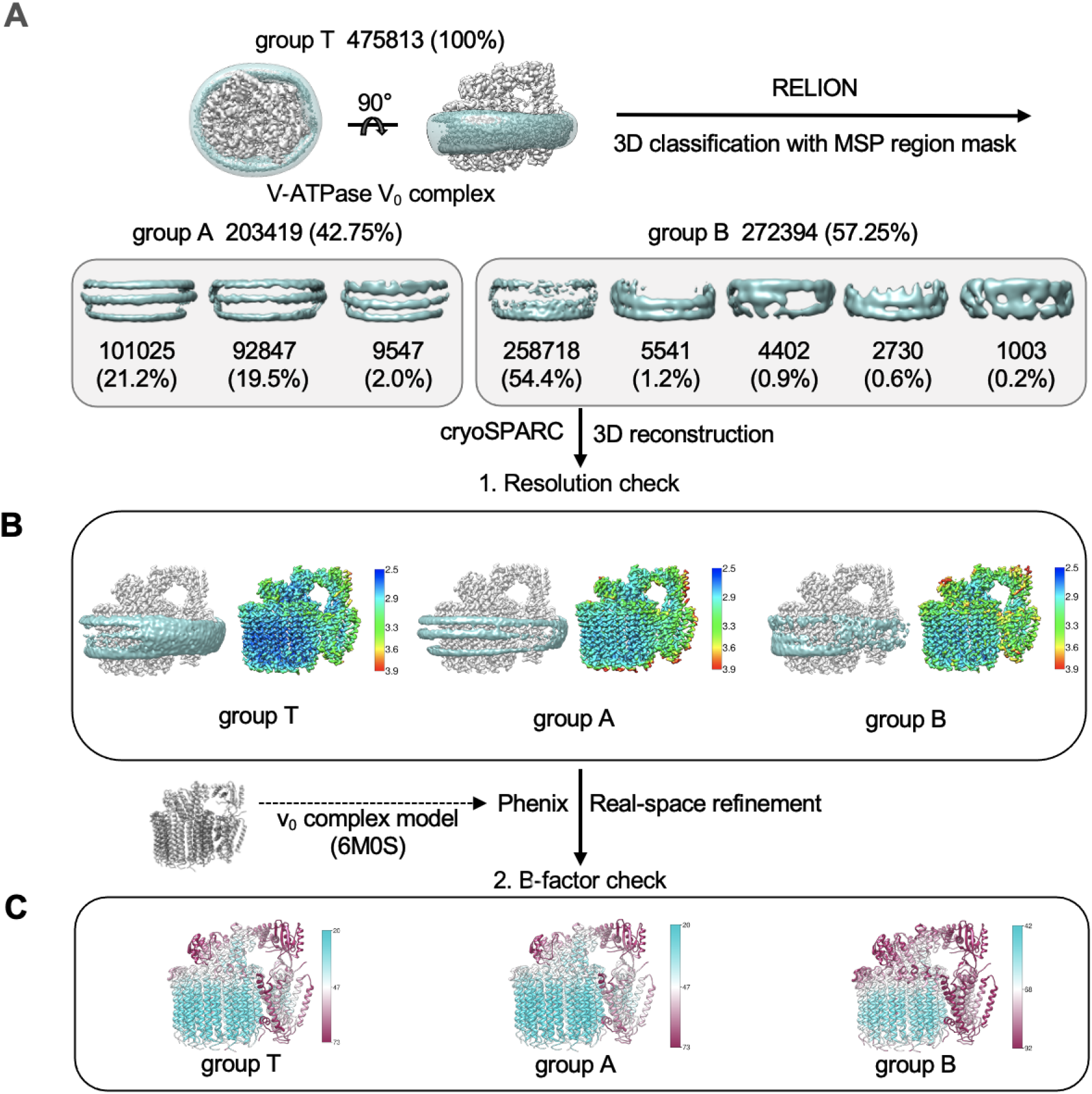
Classification and refinement of V-ATPase V_0_ complex particles by clarity of the MSP density yields differences in local resolution and B-factor values. (A) A total of 475,813 V-ATPase V_0_ complex particles were 3D-classified with a mask in the MSP region in RELION. Group A contains particles (42.75%) with distinct and well-defined MSP bands, while group B contains particles (57.25%) with disconnected and weak MSP density. Group T refers to the 3D classification performed with the entire particle set. (B) Resolution comparison of V-ATPase 3D reconstruction of each group using cryoSPARC. (*left*) Density map of the V_0_ complex (gray) and the MSP (cyan) and (*right*) local resolution colored map are shown. (C) Real-space refinement using Phenix using PDB 6M0R as a reference model yielded local B-factor values for each group.

